# No-U-Turn sampling for phylogenetic trees

**DOI:** 10.1101/2021.03.16.435623

**Authors:** Johannes Wahle

**Affiliations:** Seminar für Sprachwissenschaft, Eberhard-Karls-Universität, Tübingen, Germany

## Abstract

The inference of phylogenetic trees from sequence data has become a staple in evolutionary research. Bayesian inference of such trees is predominantly based on the Metropolis-Hastings algorithm. For high dimensional and correlated data this algorithm is known to be inefficient. There are gradient based algorithms to speed up such inference. Building on recent research which uses gradient based approaches for the inference of phylogenetic trees in a Bayesian framework, I present an algorithm which is capable of performing No-U-Turn sampling for phylogenetic trees. As an extension to Hamiltonian Monte Carlo methods, No-U-Turn sampling comes with the same benefits, such as proposing distant new states with a high acceptance probability, but eliminates the need to manually tune hyper parameters. Evaluated on real data sets, the new sampler shows that it converges faster to the target distribution. The results also indicate that a higher number of topologies are traversed during sampling by the new algorithm in comparison to traditional Markov Chain Monte Carlo approaches. This new algorithm leads to a more efficient exploration of the posterior distribution of phylogenetic tree topologies.

**Author summary:** Phylogentic trees are important for our understanding of evolutionary relationships. But even for only a small number of entities the number of possible trees is immense. In order to efficiently search through this large space and analyze the distributions of these trees, different algorithmic solutions have been proposed. A phylogenetic tree is not only defined by the way it groups the entities but also by the length of the branches. This nature of the phylogentic trees complicates the exploration of these distributions. Building on research and algorithmic ideas for non tree like data and recent developments on the mathematics of trees, I propose a new method which is able to traverse the space of possible trees more rapidly. By using a search strategy which is guided by the data and the underlying evolutionary model, the algorithm is able to better sample from the desired distribution. I am able to show that the new algorithm can indeed analyze the correct distribution but can do so with a higher efficiency than other algorithms.

## Introduction

The inference of phylogenetic trees has long been a research focus for different scholars. From the ideas of Darwin and the manually constructed trees by e.g. Ernst Häckel, to the computationally inferred variants of the *tree of life* [1], scholars have made numerous attempts to model evolutionary relationships between organisms. Apart from the biological applications, the idea of phylogenetic trees has also made its way into other disciplines such as linguistics. [2–4] This transfer of ideas dates back to the days of Darwin. [5] Bayesian approaches to infer such phylogenies use Markov Chain Monte Carlo Methods for inference. The main workhorse for prominent software solutions [6, 7] is the Metropolis-Hastings algorithm. [8, 9]

The Metropolis-Hastings algorithm is known to be rather inefficient, especially for correlated and high dimensional data. Including gradient information of the target distribution can help to solve this problem. Borrowing from physics, Hamiltonian Monte Carlo (HMC) uses this information and enriches the parameters of the model by corresponding ‘momentum’ variables to rapidly explore the target distribution. This idea already dates back to the 1980s. [10] In addition to the rapid exploration of the target distribution, the local random walk behavior of the Metropolis-Hastings algorithm can be suppressed. [11] The problem gets transformed from drawing samples to simulating Hamiltonian dynamics. For a practical implementations, the differential equations of the Hamiltonian dynamics need to be discretezied. [12]. This requires the definition of a parameter for the step-size and one for the number of steps. The successful application of a Hamiltonian Monte Carlo algorithm depends on a careful tuning of these two hyperparameters. To alleviate this problem, the No-U-Turn Sampler (NUTS) can be used. [13] This algorithm determines the optimal step-size during the burn-in phase of the experiment and adaptively sets the step-size maximizing the distance between two successive proposals.

Transferring these methods to phylogenetic research is not straightforward. The reason lies in the nature of the trees. A phylogentic tree is defined by a discrete part, the topology, and a continous part, the branch lengths. Previous research has shown that the space of trees can be formulated as a continuous space. [14] This formulation of the tree space enabled the development of a sampler based on Hamiltonian dynamics. [15] Ji et. al. [16] also propose gradient based methods for statistical phylogenetics. However, the focus of their work is the inference of branch specific evolutionary rates. Thus their project is not so much concerned with the problem of th two-part nature of the trees.

Building on the idea of the No-U-Turn sampler and the Hamiltonian style sampler for trees, I propose a No-U-Turn sampler for phylogenetic trees. The algorithm functions in a similar fashion to the classical No-U-Turn sampler and maintains the two important aspects, the tuning of the step-size and the adaptive setting of the number of steps. The important aspect is to identify the suitable distance measure. The performance of the algorithm is tested on two different data sets from the area of linguistic phylogenetics and the results of the proposed algorithm are compared against the results obtained with well established ones. It turns out that the new algorithm samples from the same distribution and that these distributions are sampled more efficiently using the new algorithm.

## Materials and Methods

### Hamiltonian Monte Carlo and the No-U-Turn Sampler

Hamiltonian dynamics provide a mathematical framework to model the dynamics of a physical system over time using the systems energy.

> In two dimensions, we can visualize the dynamics as that of a frictionless puck that slides over a surface of varying height. The state of the system consists of the position of the puck, given by a 2D vector *q*, and the *momentum* of the puck (its mass times its velocity) given by a 2D vector *p*. The *potential energy, U* (*q*), of the puck is proportional to the height of the surface at its current position, and its *kinetic energy, K*(*p*), is equal to |*p*|^2^*/*(2*m*), where *m* is the mass of the puck. [12, p. 2]

The momentum of the puck will help it to slide along a rising slope as long as the kinetic energy is above zero. At the point where the kinetic energy becomes zero it will naturally slide down again. In a non physical setting the position of the puck is equated to the variable of interest and the potential energy to the minus of the log probability density of the given variable. Only the momentum variable will be set artificially. [12] By using the Hamiltonian equations, for *d* dimensions the development of *p* and the parameters *θ* over time *t* can be determined:

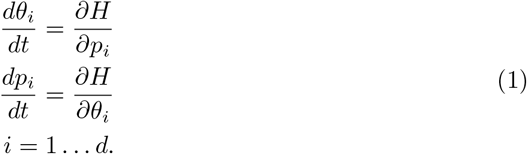

Where *H* is the Hamiltonian function taking *p* and *θ* as arguments. The function *H*(*θ, p*) can be written as

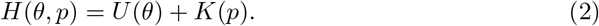

It has been shown in other places that Hamiltonian dynamics exhibit the mathematical properties which are necessary to generate valid Markov Chain Monte Carlo updates. [10, 12]

### Leapfrog approximation

For a practical implementation of Hamiltonian dynamics the continuous equations in the system need to be approximated. This can be solved by using a discretization approximation with a small stepsize *ε*. The state of the system is then computed at intervals of *ε* starting at time zero. A slight modification of Euler’s method, the *leapfrog* integrator, is commonly used as a the discretization method. Instead of doing a small step *ε* for the momentum and the position (i.e. parameter) vector in turn, the leapfrog integrator first does a half step of the momentum variables, then full step for the position variables, i.e. the variable which is sampled, and then again a half step for the momentum variable. [12]

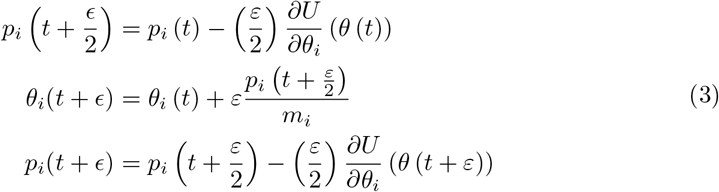

In one update step the leapfrog approximation is done for *L* steps. This describes the Hamiltonian Monte Carlo (HMC) algorithm. The discretization procedure necessarily introduces a slight error into the computation. The error depends on *ε* and *L* and can theoretically grow out of bounds depending on *L*. [17] In real world applications however, this is usually not an issue. [13] So, the leapfrog integrator fulfills all properties which are necessary for valid updates on the parameters (see Neal (2011) [12] for formal details). Algorithm 1 shows the general procedure of the Hamiltonian Monte Carlo method including the leapfrog approximation.

What remains for the user to do is to specify *L* and *ε*. In practice, setting these two parameters can be tedious. As Hoffman & Gelman (2014) [13] point out a too small *ε* will waste precious computation time while a too large *ε* will produce inaccurate samples. Similarly, if *L* is too small the HMC mirrors random walk behavior and if *L* is too large the HMC will start to loop back and retrace its steps. In some severe cases, if *L* is too large, some of the formal properties making the leapfrog method a valid sampler may even break down. [12]. Thus fine-tuning *L* and *ε* is very important.

#### Algorithm 1 HMC Pseudocode

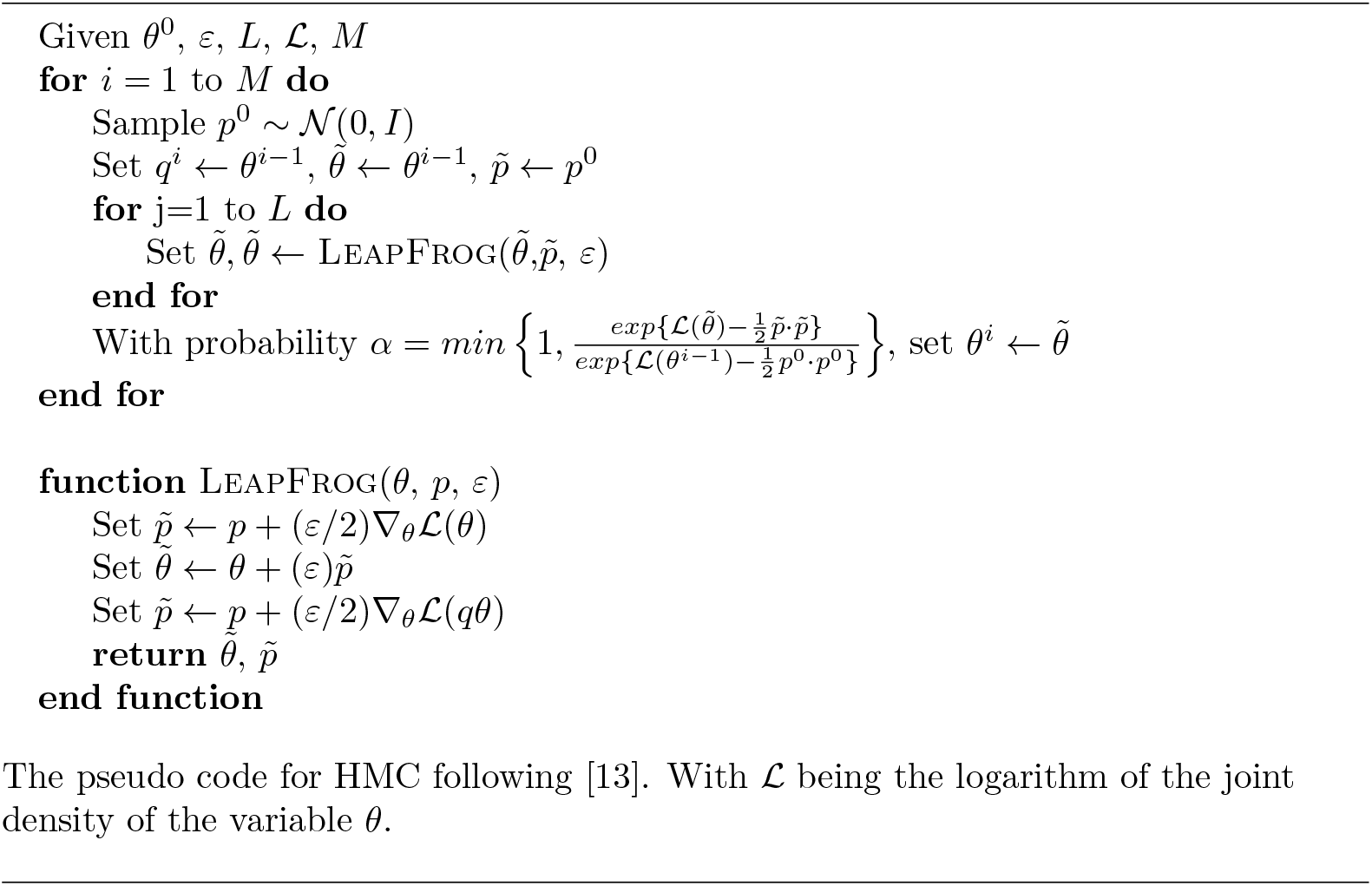

### No-U-Turn sampler

To avoid tedious and time consuming fine tuning of *L* and *ε* in several trial experiments, Hoffman & Gelman [13] developed the No-U-Turn sampler (NUTS). This sampling technique extends HMC by adaptively setting the number of steps *L* and optimizing *ε* during the burn-in phase.

NUTS uses a stochastic optimization technique to optimize *ε* during the burn-in. [13, 18] The starting value for *ε* is set by either doubling or halving *ε* until the probability of accepting an HMC update with *L* = 1 crosses 0.5.^1^ Usually the seed value for *ε* is 1. By simulating an acceptance probability of the proposed HMC moves during burn-in *ε* is altered using the dual averaging method [19] such that the overall acceptance probability approximates a given value.

In contrast to the original HMC sampling scheme where the number of leapfrog steps *L* is a fixed parameter, the number of steps in NUTS can change in each generation. Importantly, *L* is not set randomly but rather such that it maximizes the distance between successive proposals while also avoiding a loop back. Furthermore, varying the number of steps accounts for the shape of the probability density surface in the current region. The algorithm proceeds by building a set of candidate states following the slice sampling approach [20], i.e. only if a proposed state is within the slice it is added to the set. This set building procedure continues until one of two stopping criteria is reached. The two criteria which are checked, are whether the proposed states start to retrace themselves or if the sampling error introduced via the leapfrog discretization becomes too large. As soon as the extension of the candidate set is terminated a new state is randomly selected from the set. This way a maximally distant new proposal state can be ensured while avoiding sampling errors and loop back behavior. There are two points within this design of the algorithm which warrant some attention. These are the stopping rules and the formation of the set of new states.

The construction of the candidate set proceeds by simulating the Hamiltonian dynamics forward and backward in time. The final state of a forward time simulation will be called *θ*^+^ and *θ*^−^ will be the result of a backwards time simulation. Starting from the initial state, the time direction is chosen randomly. For the first iteration, the Hamiltonian dynamics are simulated for one step. For the next iteration, the dynamics are simulated for two steps and the next for four steps. So every new iteration doubles the number of leapfrog steps taken. This procedure implicitly builds a balanced binary tree with the leaves corresponding to a pair of position-momentum vectors *p* and *q*. [13] More formally, let *C* be the set of candidate states and let *B* ⊇ *C* be the set of states a leapfrog integrator visits during an iteration. Hoffmann & Gelman [13] show that as long as the initial state of the current generation is an element of *C* and for all other elements of *C* it is the case that they are in the current slice and the stopping criteria are not met, sampling a random element from *C* is a valid update. Building up *C* in this way is only valid if the method doing so preserves volume.^2^ Neal (2011) [12] show that the leapfrog integrator does preserve volume, so it can be used to propose new states for *C*. While the algorithm for building candidate sets and sampling from it fulfills all necessary conditions for being a valid sampler and also proposes distant states from the starting state, Hofmann & Gelman [13] modify their initial algorithm slightly to improve its memory efficiency and also let it do larger jumps on average. The modifications include an earlier breakout of the simulation procedure if one of the stopping criteria is met, and a more efficient sampling from *C* which prevents the storing of all states in memory.

In comparison to the intricate mechanism of proposing new candidate states, the stopping rules of NUTS are rather straightforward. Recall the two tests, which stop further simulation of Hamiltonian dynamics. The first checks for sampling inaccuracies and the second for an indication of retracing of states. Formulating the first condition is rather uncomplicated. Hofmann & Gelman [13] suggest to compare the Hamiltonian to some constant Δ_*max*_ and check if the loss of energy is larger than this constant. The other condition, which checks whether the simulation of the Hamiltonian dynamics would suggest that a retracing of states is going to happen. This condition is checked via simulating the dynamics for another infinitesimal amount forward and backward. If this simulation would reduce the distance between the new states, the simulation is stopped.^3^

### Hamiltonian Dynamics in the Tree Space

Hamiltonian Monte Carlo methods are defined on a Euclidean parameter space. Using Hamiltonian Monte Carlo to sample phylogenetic trees is not straightforward due to their double nature. They are defined by two parts, a discrete graph defining the topology and a set of non-negative numbers specifying the length of each edge. Importantly, there exists the formulation of a continuous space which models all phylogenetic trees over a fixed set of leaves. [14] Building on this work, Dinh et. al. [15] proposed a Hamiltonian Monte Carlo sampler for trees.

The set of all binary phylogenetic trees with *N* leaves will be called *T*_*N*_. An element of the set *T*_*N*_ is a pair (*τ, q*), where *τ* defines the topology and each *q*_*e*_ ∈ *q* is the length of a particular edge. Let Γ be the set of all *τ*. The size of *q* equals the number of edges of *τ*. *Pendant* edges are those incident to leaves. Any other edge is called *internal*. Consider a function *f* (*τ*) → *q* which maps the length of each edge of *τ* to a specific component of *q*. For a particular tree topology *τ* there is a vector 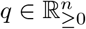, where *n* is the dimension of *q*. The individual branch lengths of pendant edges are bounded by 0 for technical reasons.^4^ [15] For each topology *τ* of the trees in *T*_*N*_ there is a specific space 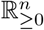 associated with it. For the specific case where for each topology *τ* the value of each entry of *q* is 0 all trees are indistinguishable. This point is called the origin of the space. These different spaces 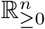 are associated with different topologies. These orthants are part of the same orthant complex. For *N* leaves there are (2*N*− 3)!! such orthants. The set Γ can be used to index the different orthants. This formulation neatly gives rise to an intuitive neighboring relation between trees embedded in this space. Imagine reducing the length of an internal edge of a particular tree to zero. In this case, the two nodes connected by this edge, would collapse into one vertex with degree four. By expanding the resulting degree four vertex in an edge and two vertices with degree three, a new tree topology can be created. This operation is also known as a nearest neighbor interchange (NNI) operation (see Fig. 1). [15, 24] By using this relation, orthants are called neighbors if there exists one NNI operation which can transform the one topology into the other. Neighboring orthants share a so called *facet*. The facet of an orthant in this case is a lower dimensional vector of branch lengths. The tree in the middle of Fig. 1 has a branch length vector with lower dimensionality than the left or right one.

**Fig 1.**
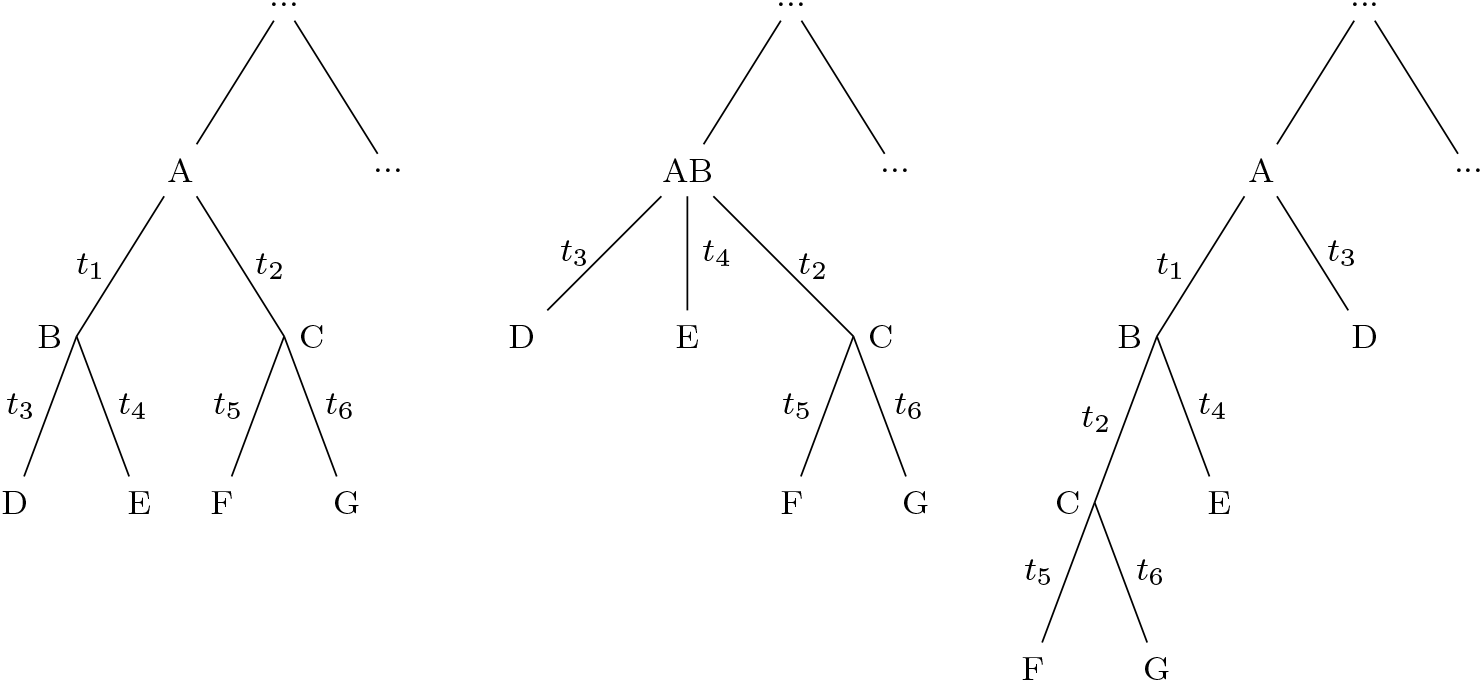
Example of a NNI move. The rightmost tree can be generated from the leftmost by a NNI move. The tree in the middle displays the situation where the length of the branch *t*_1_ was reduced to 0 from which the new tree can be generated.

Importantly, Billera, Holmes & Vogtmann [14] show that this space of trees has certain properties such that a *geodesic* between two trees exists. Thus, the distance between two trees in this space is properly defined. Amenta et. al. [25] show how the lower and upper bound on this distance can be calculated in linear time and Owen (2008) [26] presents an exact algorithm. After the authors fo the original paper, this space of trees is called a Billera-Holmes-Vogtmann space.

Calculating the likelihood *L* (*τ, q*) for a phylogenetic tree (*τ, q*) ∈ *T*_*N*_, can be done using Felsensteins Pruning algorithm. [27] Given a set of *N* observed sequences of length *S* with characters from an alphabet Ω the likelihood can be computed as follows. Let 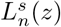 be the likelihood of node *n* at site *s* ∈ *S* being in state *z*

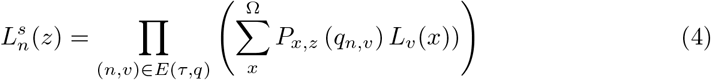

where *P*_*x,y*_ (*t*) is the probability of transitioning from state *y* to state *x* in time *t* and *E* (*τ, q*) is the set of all edges of the tree, such that (*u, v*) ∈ *E* (*τ, q*), where *u* is the mother node of *v*. At the leaves of the tree, *L*_*n*_(*s*) is 1 if *s* is the observed character and 0 otherwise. Let *r* be the root node of the tree and *π* (*a*) be the equilibrium frequency of character *a*, then the likelihood can be computed as follows.

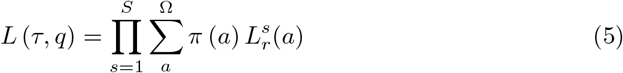

With the formulation of the space of trees and the description of the likelihood a Hamiltonian Monte Carlo sampler for (phylogenetic) trees can be defined. To ensure that there is a one-to-one correspondence between the attributes of *q, f* (*τ*) needs to be consistent. Consistency can be defined as follows: For two topologies *τ, τ* ^′^ ∈ Γ let *f* (*τ*) = *f* (*τ* ^′^) at the boundaries of the orthants of *τ* and *τ* ^′^.

Since the space within one orthant is continuous, the traditional HMC procedure can be applied. It only becomes problematic when the boundary of an orthant is reached which means that one particular element of the branch length vector becomes zero or even negative. Through the way the space of trees is defined, it is possible to interpret the scenario when an element of *q* becomes zero. In the case of a rooted binary tree there are three topologies, two generated via an NNI operation and the original, which share this particular facet. The sampler will pick one of these three orthants at random and will continue the simulation in the newly selected orthant. The momentum corresponding to that particular attribute will then be negated. The Hamiltonian remains unaffected by using the described technique. [15] This defines the “leap-prog” integrator, the probabilistic counterpart to the standard leapfrog integrator. If in a current update step multiple elements of *q* hit zero, the random selection process will be done for each of this elements. Thus, it is possible to visit multiple tree topologies in just one leap-prog step.

As an extension to this simple algorithm Dinh et. al. [15] note that via the discontinuity of the space of trees, the potential energy function on *τ* and *q* which is the likelihood function, is not differentiable on the whole space and may thus lead to a loss of accuracy which can not be neglected. To circumvent this problem the authors propose to use the following smooth approximation for (*τ, q*) [15]

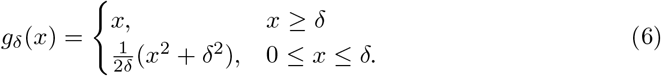

This smoothing function is applied to each element of *q* and ensures that the gradient vanishes when the element approaches 0, thus the derivative to be continuous across orthants. As a trade-off however, the likelihood function is not continuous anymore across orthants. To alleviate this problem the state resulting from a topology move is accepted using the original Hamiltonian. As a side effect, a particular choice of *δ* also results in a lower bound for the individual branch lengths. The minimum of *g*_*δ*_(*x*) is at 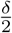, which will effectively function as the lower bound. It can be shown that the Hamiltonian dynamics defined above uphold the reversibility, volume preservation and k-accessibility properties of traditional Hamilitonian dynamics. [15]

### No-U-Turn Sampling in the Tree Space

There are two aspects of NUTS which need to be adapted to define the pyhlogenetic No-U-Turn sampler (p-NUTS). The first part is to provide a HMC sampler in the space of trees and the second is to define an appropriate stopping rule for the NUTS tree building procedure.

Replacing the leapfrog integrator in the No-U-Turn sampler with the leap-prog integrator described by Dinh et. al [15] is straightforward. Since the leap-prog integrator preserves volume and is reversible as the leapfrog integrator, the replacement does not impact the target distribution. [15] Thus, using the leap-prog integrator instead of of the traditional leapfrog method maintains the theoretical properties of original No-U-Turn sampler. Since Dinh et. al. [15] report a superior performance for the leap-prog integrator which uses the smoothed branch lengths, this variant will also be used for p-NUTS.

The second aspect which needs to be defined is an appropriate stopping rule for the number of steps. For NUTS, this procedure is stopped if an infinitesimal change in either direction would cause the states at the left- and rightmost part of the search tree to start moving closer together. For p-NUTS, this means a comparison of trees is necessary. The fact that a geodesic between any two trees in the Billera-Holmes-Vogtman space exists can be leveraged here. However, calculating the actual distance becomes infeasible with an increasing number of leaves. [25] Instead, the lower and upper bound on the geodesic are used. These bounds will function as proxies for the actual distances. The algorithms to calculate these bounds are in linear time, while the best algorithms for the actual distance are exponential in the number of leaves. [25] Using the lower and upper bounds as distance proxies results in the following procedure for checking the No-U-Turn behavior.

1. simulate one leap-prog step for *θ*^−^ and *θ*^+^ resulting in *θ*^−′^ and *θ*^+′^ respectively
2. Calculate the lower bound on the distance of *θ*^+′^ and *θ*^−′^, *D*_*l*_ and the upper bound *D*_*u*_ for *θ*^+^ and *θ*^−^
  a. if *D*_*l*_ *< D*_*u*_ terminate the sampling step
  b. else continue sampling

As for standard NUTS the simulation is also aborted if the error in the simulation becomes too large. The same rule as described above will be used. This new stopping rule also leaves the target distribution intact since the target distribution is not affected by any of the operations in the No-U-Turn measure. The difference to standard NUTS lies in the more complex evaluation of *θ*^−′^ and *θ*^+′^. While the standard algorithm only requires the computation of two inner products, the p-NUTS sampler requires the evaluation of an entire leap-prog step. However, as soon as the height of the binary search tree *j* becomes larger than 2, computing two leap-prog steps is cheaper than computing another 2^(*j*+1)^ new leap-prog steps. This concludes the definition of the stopping rule for p-NUTS.

The second part of the No-U-Turn sampler is the automated tuning of the step-size. Hofmann & Gelman [13] propose to adapt the step-size with respect to a target acceptance probability. For the p-NUTS sampler there is a second target quantity which should influence the optimal step-size. This second quantity is the number of NNI moves. As strikingly shown by Whidden & Matsen [28], the highest posterior probability trees for a given experiment do not need to have a small distance between each other. In order to bridge the gaps between such very different tree topologies a high number of NNI moves might be necessary. Thus, optimizing the step-size for the number of NNI moves as well is necessary. Let *H*^*NUT S*^ be the estimate of the average acceptance probability during the last iteration of the tree building process as defined by Hofmann & Gelman [13]. Then *H*_*t*_ = *δ* − *H*^*NUTS*^ with *δ* as the target average acceptance probability is a statistic describing the behavior of the MCMC which can be used as a condition for the dual averaging scheme. In order to incorporate the number of NNI moves, let *η* be the target number of NNI moves and *H*^*PN*^ be the estimate of the average number of NNI moves during the last iteration of the tree building process. Let *H*_*p*_ be the statistics describing the behavior of the MCMC regarding the number of NNI moves.

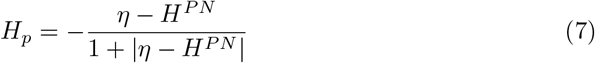

This statistic is nonincreasing in the step-size, which is a necessary condition, and can thus be used to describe the behaviour of the MCMC as well. In order to combine *H*_*t*_ and *H*_*p*_, their average is computed as the statistics describing the P-NUTS sampler.^5^ The remainder of the dual averaging paradigm described by Hofmann & Gelman remains unchanged.

### Data

In order to assess the performance of the p-NUTS sampler some empirical tests were carried out. Using two different datasets from the supplementary material of Jäger (2018) [29], the new p-NUTS method and MrBayes [6] are used to infer posterior distributions over the phylogenetic trees. Table 1 gives an overview over the relevant statistics of the data sets that were used to test p-NUTS. The data show different relations between the number of sites and the number of languages.

**Table 1.**
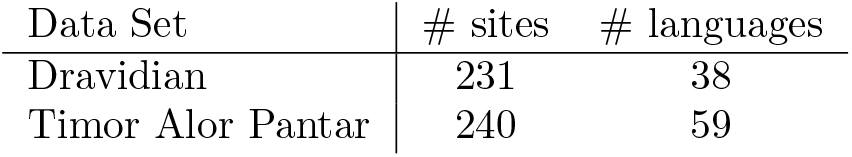
Overview of the data sets used to test p-NUTS.

| Data Set | # sites | # languages |
| --- | --- | --- |
| Dravidian | 231 | 38 |
| Timor Alor Pantar | 240 | 59 |

## Results

The first step in assessing the correctness of the new sampler is to identify if it samples from the correct distribution, the second step will be to assess the speed of convergence for the new p-NUTS algorithm. Assuming that MrBayes indeed samples from the correct distribution, MrBayes was run for 10^6^ generations. Since the aim is not to infer a perfect phylogenetic model for the three different data sets, but to test how well NUTS can sample tree structures, the only parameters inferred are the trees and the equilibrium frequency of the presence/absence of the binary coding character.

Table 2 provides a summary of the results of the different analyses.^6^ Overall the data in the table shows that p-NUTS and MrBayes converge to similar posterior distributions. This statement is also confirmed by comparing the distributions of the *Generalized Quartet Distances* [30] of the trees from the respective posterior samples to the gold standard trees (c.f. Fig. 2).

**Table 2.** Results from the MCMC runs for MrBayes and p-NUTS. Results from the MCMC runs for MrBayes and p-NUTS. *π* is the equilibrium frequency of the presence of the character, TL the tree length, LL the log-likelihood of the model. The mean values of the posterior distributions are reported here, the values in the brackets are the standard deviations.

|  | Dravidian |  |  |
| --- | --- | --- | --- |
| | $\pi$ | TL | LL |
| MrBayes | 0.781($\pm 0.014$) | 2.798($\pm 0.171$) | -1604.046($\pm 7.18$) |
| p-NUTS | 0.777($\pm 0.015$) | 3.075($\pm 0.223$) | -1621.426($\pm 11.58$) |
|  | Timor Alor Pantar |  |  |
| | $\pi$ | TL | LL |
| MrBayes | 0.831($\pm 0.010$) | 3.535($\pm 0.20$) | -2263.246($\pm 9.14$) |
| p-NUTS | 0.826($\pm 0.013$) | 3.803($\pm 0.258$) | -2282.459($\pm 39.50$) |

**Fig 2.**
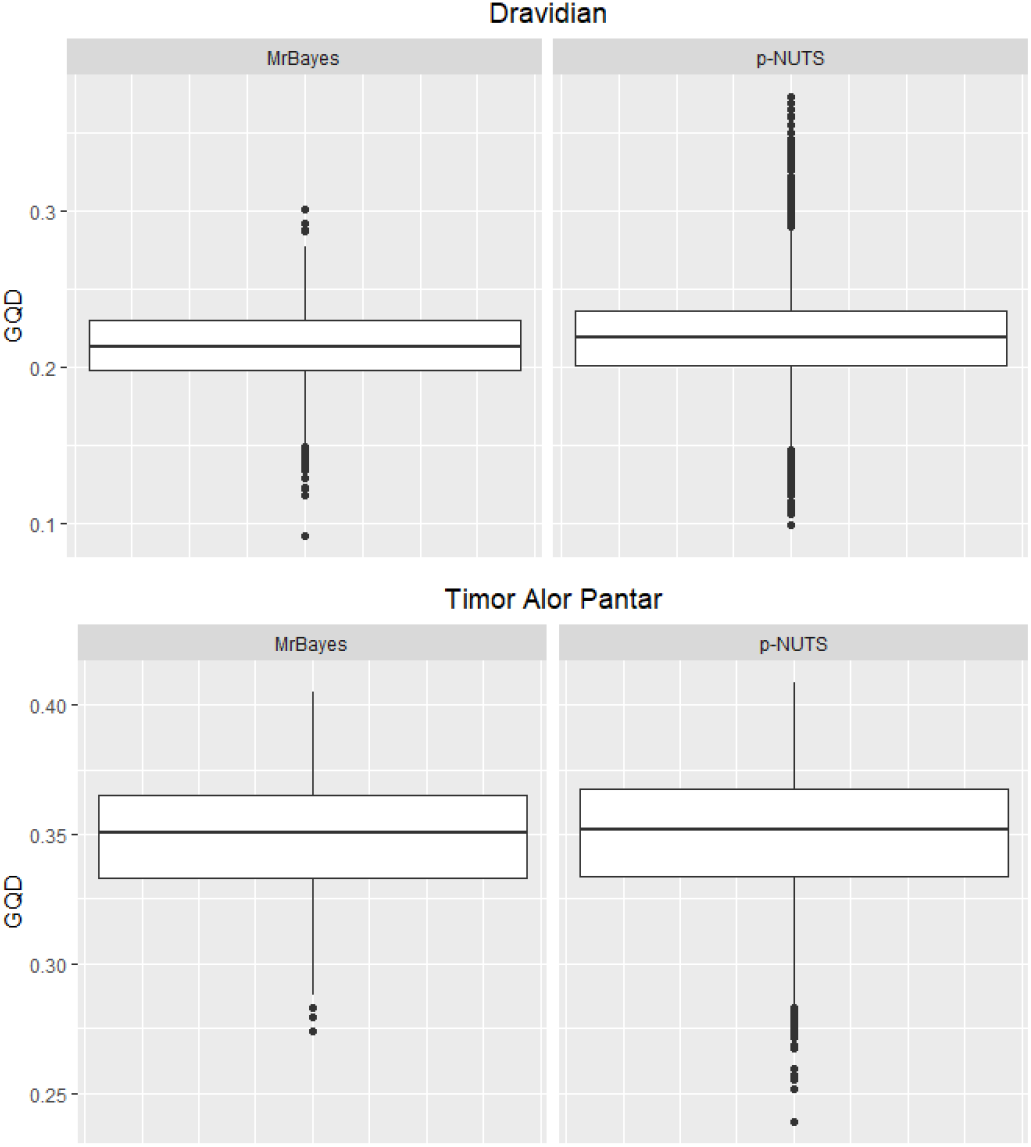
Distribution of GQDs. Distribution of Generalized Quartet Distances (GQD) between the gold standard tree and the trees in the posterior sample of PNUTS and MrBayes.

The next step in evaluating p-NUTS is assessing its speed of convergence. In order to do so, I will focus only on the tree length. Figure 3 shows the development of the shrink factor against the number of iterations in the MCMC runs. As can be seen from the plot, the shrink factor approaches 1 for each of the p-NUTS runs rapidly. This is somewhat expected as already the original NUTS sampler is able to produce nearly independent samples in every generation. This behavior is mirrored by p-NUTS. This result is in line with the observations made for the probabilistic path sampler. [15]

**Fig 3.**
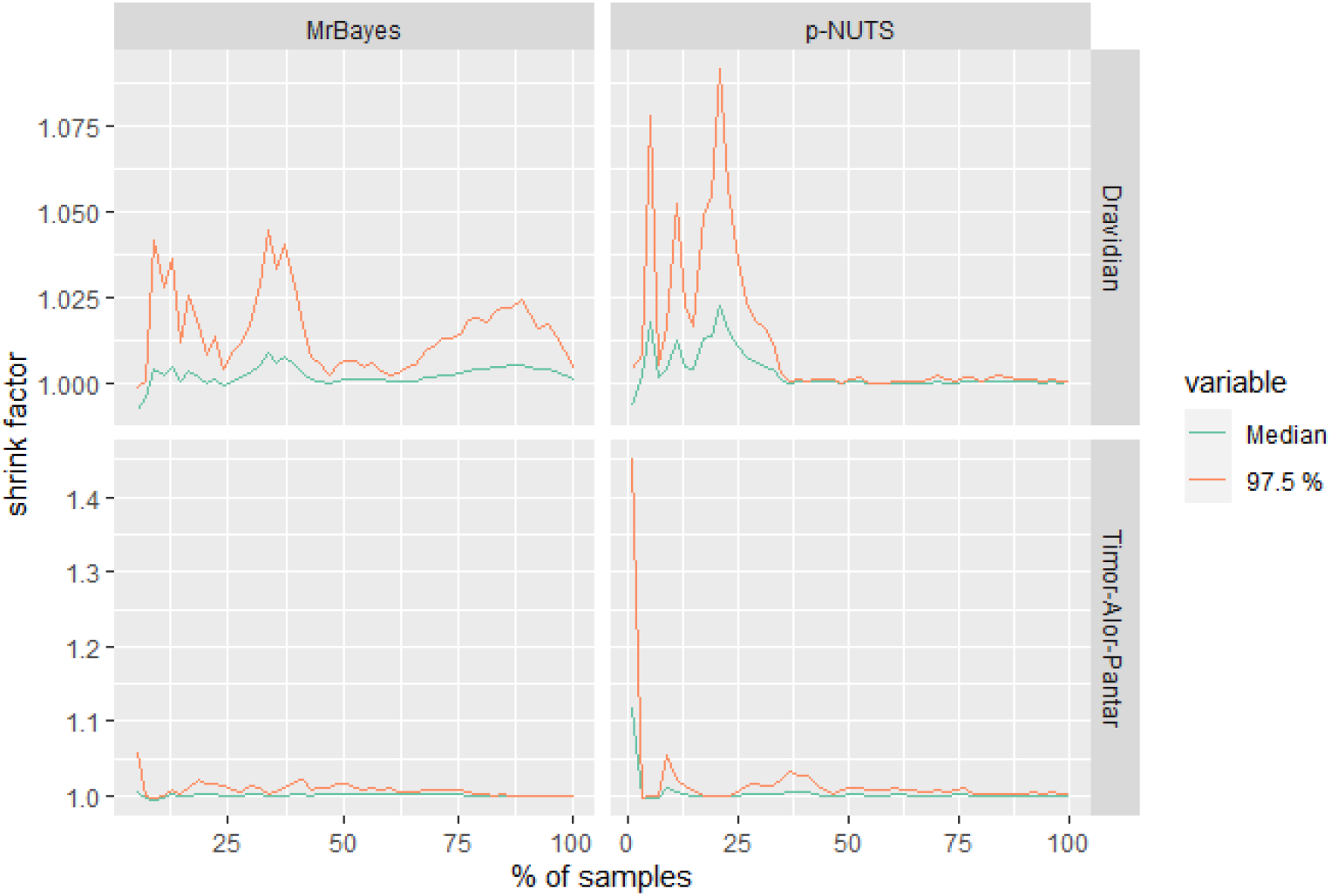
Shrink factor against iterations. This plot shows the development of the shrink factor against the percentage of sampled iterations for each of the data sets. [31, 32]

Another statistics supports the effectiveness of the p-NUTS sampler. The numbers in table 3 list the average number of NNI moves per generation for the p-NUTS sampler.

**Table 3.**
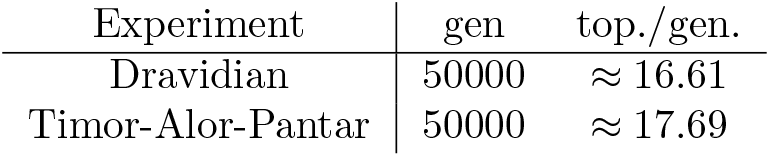
Topologies per generation for p-NUTS. The table lists the number of generations for each experiment and average number of (not necessarily different) the topologies visited per generation for the p-NUTS sampler.

| Experiment | gen | top./gen. |
| --- | --- | --- |
| Dravidian | 50000 | $\approx 16.61$ |
| Timor-Alor-Pantar | 50000 | $\approx 17.69$ |

Although these are not necessarily different topologies, the p-NUTS sampler still explores the tree space much faster than a traditional MCMC method. The number of NNI moves for the experiments on the Dravidian and Timor-Alor-Pantar data set exceeds the number of NNI moves performed during 10^6^ generation in the MrBayes experiments.

### Computational Performance

Although p-NUTS needs a considerably smaller amount of generations to sample from the target distribution, the current algorithm is not yet up to speed or even faster than MrBayes. By the design of the algorithm, the likelihood function needs to be evaluated for each proposed tree move. So for all the experiments, the number of tree topologies visited per generation by p-NUTS is the minimal number of times the likelihood function is evaluated per generation. As can be inferred from table 3, this number is always considerably larger than the number of generations in the experiments done with MrBayes. In addition to calculating the likelihood function, p-NUTS requires several evaluations of the gradient, which adds further computational burden. Implemented in the *Julia* programming language [33], the current version of p-NUTS can calculate 50000 generations in one and a half hours for the Dravidian data set, which amounts to roughly 555 generations per minute.^7^ In addition to the costly p-NUTS sampler for the phylogenetic tree, the equilibrium frequencies are sampled using a slice sampler. The slice sampler may also require multiple evaluations of the likelihood function per generation. On the same machine, MrBayes can finish the calculations in a matter of minutes.^8^

## Discussion

The algorithm presented in this paper can be used to infer a posterior distribution of phylogoentic trees based on a gradient based sampling scheme. Ji et. al. [16] also propose gradient based methods for statistical phylogenetics. ^9^ However, the focus in their paper is the inference of branch specific evolutionary rates. Although based on a similar algorithmic idea, the focus of the two projects is clearly different. In addition to their results, Ji et. al. [16] show that the gradient calculation can be done in linear time of the number of sampled sequences. Experiments were conducted with both an automatic differentiation library calculating the gradient [35] and a static implementation of the gradient. Both versions were not able to bring p-NUTS up to speed with MrBayes in terms of mere run-time. Furthermore, it is desirable to find a way to eliminate the spurious calculations of the likelihood function in while checking the stopping criterion, which in its current state requires the simulation of a full leap-prog step.

The Stan platform [36] for statistical computation proposes further algorithmic improvements for HMC based samplers. These include other ways of generating the kinetic energy function, which can lead to improved convergence and in the presence of correlated parameters, or improved sampling strategies in the final step of the NUTS algorithm. These options can help to improve the performance of the p-NUTS sampler. However, for further run time improvement of the p-NUTS algorithm the very first issue seems to be the most important next step.

## Conclusion

The paper proposes an algorithm which is capable of performing No-U-Turn sampling for phylogenetic trees. Despite the complicated structure of the space of trees, Billera, Holmes & Vogtmann [14] were able to define the space of trees in such a way that Hamiltonian Monte Carlo sampling is possible. The leap-prog approach for sampling phylogenetic trees, brought forward by Dinh et. al. [15], lends itself naturally to build an algorithm for phylogenetic No-U-Turn sampling. The results show that indeed better mixing and faster convergence is achieved by using p-NUTS to sample phylogenetic trees.

## Footnotes

1 A HMC update with *L* = 1 is also known as a Langevin proposal.

2 See Betancourt (2018) [21] for a conceptual overview over Hamiltonian dynamics and volume preservation.

3 Betancourt [22, 23] presents a generalization the No-U-Turn measure.

4 Furthermore, there seems to be no intuitive interpretation of a tree with negative branch length in phylogenetic applications.

5 Care has to be taken in setting *η*. If the value is too large the step-size parameter might get too large and the algorithm becomes unstable (see Neal 2011 [12]).

6 It has to be kept in mind that p-NUTS has a lower bound on the branch lengths through the usage of the smoothing parameter *δ*. This may lead to slightly different estimates in the exact branch lengths of the tree.

7 Times are measured on an *AMD Ryzen 9 3900X CPU* with 12 cores and 2.9 GHz. Although the particular version of p-NUTS used only 5 cores maximally.

8 On a side note, MrBayes is implemented in the C programming language. To compare p-NUTS and MrBayes on equal footing, both approaches would need to be written in the same programming language. This however, would likely not change the quality of the runtime comparison, because of the increased complexity of p-NUTS. See Hoffman & Gelman (2014) [13] for a short discussion about the run time of Hamiltonian Monte Carlo methods.

9 See also Kenney & Hu [34] for derivations of the gradient of the likelihood function.

